# Amyloid β accelerates age-related proteome-wide protein insolubility

**DOI:** 10.1101/2023.07.13.548937

**Authors:** Edward Anderton, Manish Chamoli, Dipa Bhaumik, Christina D. King, Xueshu Xie, Anna Foulger, Julie K. Andersen, Birgit Schilling, Gordon J. Lithgow

**Affiliations:** The Buck Institute for Research on Aging, 8001 Redwood Blvd, Novato, CA 94945; USC Leonard Davis School of Gerontology, University of Southern California, 3715 McClintock Ave., Los Angeles, CA 90191

## Abstract

Loss of proteostasis is a highly conserved feature of aging across model organisms and typically results in the accumulation of insoluble protein aggregates. Protein insolubility is a central feature of major age-related neurodegenerative diseases, including Alzheimer’s Disease (AD), where hundreds of insoluble proteins associate with aggregated amyloid beta (Aβ) in senile plaques. Moreover, proteins that become insoluble during aging in model organisms are capable of accelerating Aβ aggregation in vitro. Despite the connection between aging and AD risk, therapeutic approaches to date have overlooked aging-driven protein insolubility as a contributory factor. Here, using an unbiased proteomics approach, we questioned the relationship between Aβ and age-related protein insolubility. We demonstrate that Aβ expression drives proteome-wide protein insolubility in C. elegans and this insoluble proteome closely resembles the insoluble proteome driven by normal aging, suggesting the possibility of a vicious feedforward cycle of aggregation in the context of AD. Importantly, using human genome-wide association studies (GWAS), we show that the CIP is replete with biological processes implicated not only in neurodegenerative diseases but also across a broad array of chronic, age-related diseases (CARDs). This provides suggestive evidence that age-related loss of proteostasis could play a role in general CARD risk. Finally, we show that the CIP is enriched with proteins that modulate the toxic effects of Aβ and that the gut-derived metabolite, Urolithin A, relieves Aβ toxicity, supporting its use in clinical trials for dementia and other age-related diseases.

## Introduction

Insoluble protein aggregates accumulate during the normal aging process across eukaryote species, from yeast to mice^1–9^. In *Caenorhabditis elegans*, insoluble protein accumulation is strongly associated with lifespan^4^. Chemical compounds that slow the accumulation of insoluble proteins extend lifespan^10^; whereas a diet rich in iron accelerates protein insolubility and shortens lifespan^11^. In humans, protein insolubility and dysregulation of proteostasis are central phenomena of all major, age-related neurodegenerative diseases, despite the canonical protein associated with aggregation being different in each case. Genetic, biochemical, and animal model data all support a central role for Aβ, and in particular the Aβ_1–42_ peptide, in the formation of aggregates and neurodegeneration in Alzheimer’s Disease (AD)^12–21^. However, senile plaques and neurofibrillary tangles contain hundreds of insoluble proteins, in addition to Aβ and Tau, and many of these same proteins have been found to aggregate during normal aging in *C. elegans*^4,5,9,22,23^. Intriguingly, insoluble protein extracts, specifically from old animals, significantly accelerate the aggregation of Aβ *in vitro*, suggesting a biophysical relationship between aging insoluble proteome and Aβ aggregation^24^. While aging is the dominant risk factor for AD, therapeutic approaches have thus far failed to consider age-related generalized protein insolubility as a contributory factor^25–28^. We questioned the impact of Aβ accumulation on the insolubility of proteins that typically insolubilize during aging? Could it be the case that age-dependent insoluble proteins interact with Aβ in a destructive feedforward cycle, leading to an acceleration of protein insolubility in AD? We propose that by understanding which proteins are affected in this way we could help uncover a novel mechanism to prevent Aβ toxicity.

Here, using an unbiased proteomic approach, we tested the effect of Aβ expression on protein insolubility using a well-established *C. elegans* model ^29^. We demonstrate that Aβ expression drives a dramatic increase in proteome-wide protein insolubility, and that this insoluble proteome is highly similar to the insoluble proteome that forms due to normal aging. Having identified a highly vulnerable insoluble sub-proteome we term it the Core Insoluble Proteome (CIP). By analyzing human genome-wide association studies (GWAS), we show that the CIP is filled with biological processes implicated across not only neurodegenerative diseases but also diverse chronic, age-related diseases (CARDs), providing suggestive evidence that protein insolubility could play a role in general CARD risk. Finally, we show that the CIP can be targeted genetically or pharmacologically to modify Aβ toxicity. Taken together, our findings provide insights into Aβ toxicity in AD and highlight the importance of considering the insoluble proteome in not only AD but also in other age-related diseases.

## Results

### Aβ expression drives proteome-wide protein insolubility resembling normal aging

To interrogate the effect of Aβ on proteostasis we utilized the *C. elegans* strain, GMC101, which expresses the human, pro-aggregating, and pathogenic Aβ_1–42_ peptide in muscle tissue, here referred to as simply Aβ^18,30^. When cultured at 20ºC, worms express Aβ at low levels and when shifted 25ºC overexpress Aβ. After 24 hours of exposure, the majority of worms paralyze^30^. We allowed worms to develop in the absence of Aβ expression and then moved them to 25ºC at the beginning of adulthood. Worms were then collected after 24 hours of exposure to Aβ when most worms were paralyzed. We extracted insoluble aggregated proteins by serial washing worm lysates with 1% Sodium Dodecyl Sulfate (SDS) buffer. We used mass spectrometry and data independent acquisitions (DIA) to identify and quantify proteins, as previously described^29,31,32^ (Fig.1A). Background genotype-control worms (CL2122) lacking the Aβ transgene were cultured and processed in parallel for comparison (Fig.1A).

**Figure 1:**
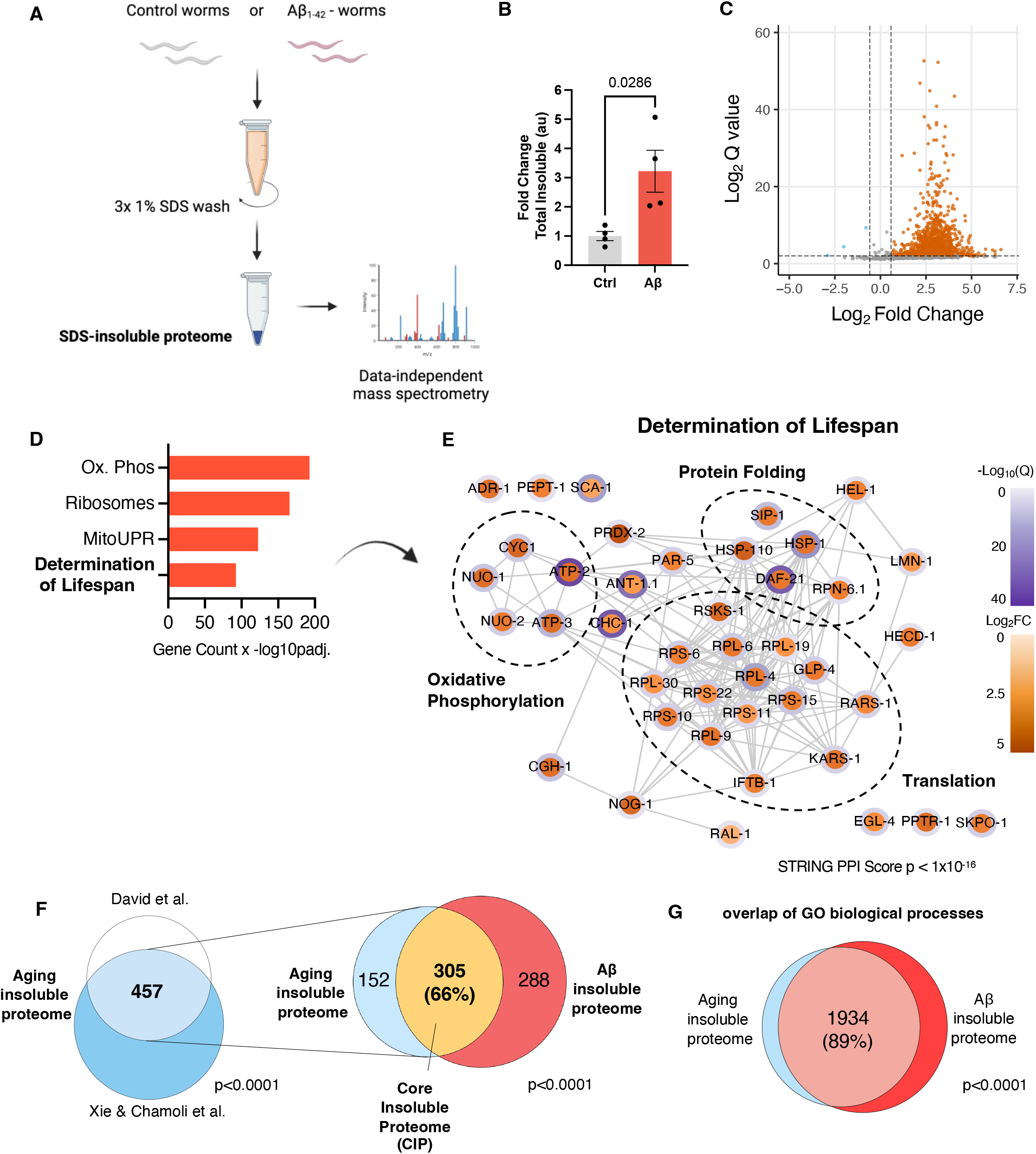
Aβ drives proteome-wide protein insolubility that resembles normal aging. **A**. Schematic of experimental procedure. **B**. Total insoluble protein Intensity in worms expressing Aβ vs genotype control strain CL2122, all values normalized to the average of the control in each experiment n = 4, error bars = SEM, Mann-Whitney Test. **C**. Representative volcano plot from one experiment Absolute Log_2_(Fold Change)>0.58, Q <0.01 **D**. Barplot of selected enriched GO terms, KEGG pathways and Wikipathways in the Aβ insoluble proteome, Benjamini Hochberg FDR <5% (see supplementary tables for complete list) **E**. STRING protein-protein interaction network of insoluble proteins with annotations for “Determination of Lifespan”, average fold change and Q values across both experiments, STRING p < 1×10^−16. **F**^. left: **F**. Overlap of proteins enriched in the insoluble proteome of old *C. elegans* from two publicly available datasets^43,49^ p<0.0001 Fischer’s Exact Test. right: Overlap of aging insoluble proteome with Aβ insoluble proteome, p<0.0001 Fischer’s Exact Test. **G**. Overlap of all GO annotations from the aging insoluble proteome and Aβ insoluble proteome, p<0.0001 Fischer’s Exact Test.

Aβ expression in young adult animals caused a robust increase in the total amount of insoluble protein (Fig.1 B&C). We identified and quantified peptides representing 1704 proteins in the insoluble fraction across four biological replicates, of which 593 proteins robustly increased due to Aβ expression across independent experiments (Fig. S1A). We observed a proteome-wide increase in insolubility, impacting a wide array of essential functions (Fig S1B). Cytoplasmic and organelle-specific heat shock proteins (HSPs) from across major organelles were enriched in the insoluble fraction (Fig S1B). Several proteins involved in maintaining proteostasis also became insoluble such as the proteasome regulatory ‘lid’ complex, the TriC chaperonin complex, and key lysosomal proteins (Fig S1B). Furthermore, we observed an increase in protein insolubility of almost every ribosomal subunit and translation accessory factors. The Aβ insoluble proteome was highly enriched with mitochondrial proteins (109/593); particularly those involved in the ETC and TCA cycle (Fig. S1B, Supp. Table 1). Consistent with this, the mitochondrial unfolded protein response (mitoUPR) chaperone hsp-6 increased 9.2-fold in the insoluble fraction and GO enrichment analysis revealed mitoUPR was amongst the most significantly enriched pathways (Fig. 1D, Supp. Table 1 & 2). Strikingly, 43 of the 88 ETC proteins became insoluble due Aβ expression (ETC complexes detailed in Supp. Table 3). We noted that nuclear-encoded ETC complexes, were particularly vulnerable to insolubility (Fig. S1B) but this was not accompanied by insolubility of the mitochondrial DNA-encoded ETC complex proteins: only one mtDNA-encoded, complex IV protein became insoluble (Supp. Table 3). The skewed representation of nuclear-encoded subunits could be explained by the fact that we also observed a significant increase in insolubility of mitochondrial outer membrane proteins responsible for importing nuclear encoded ETC proteins. All major proteins of the TOM complex: TOM-20, TOM-22, TOM-40, and TOM-70 became insoluble (Supp. Table 1). Single nucleotide polymorphisms (SNPs) in several TOM complex genes have been linked with AD risk, most prominently *TOMM40* ^33–35^ (Fig. S1B). We also identified GOP-3, the orthologue of the outer membrane complex protein SAMM50^36^, which is required for threading of mitochondrial β-barrel proteins into the outer and inner membrane bilayers^37^, along with several small molecule transporters: VDAC-1, ANT-1.1 and MTCH-1 (Fig S1B, Supp. Table 1). We questioned if many of the proteins in the Aβ-driven insoluble proteome have been shown to interact or co-aggregate with Aβ. To assess this, we compared laser microdissection proteomics data from human AD senile plaques with the Aβ-driven insoluble proteome and found a highly significant overlap (Fig S1C, Supp. Table 4). Specifically, ∼1/3 of the proteins which co-aggregate with Aβ in plaques also become insoluble due to Aβ expression in *C. elegans*, almost all of which are intracellular proteins (Fig. S1C & S1D, Supp. Table 4)^38^. Similarly, we find that worm orthologues for 15 of the 28 proteins identified to reproducibly interact with Aβ proteins in transfected cells, were enriched in the insoluble proteome after Aβ expression in *C. elegans*^39^ (Supp. Table 4).

Gene Ontology (GO) and pathway enrichment analysis revealed that oxidative phosphorylation (Ox. Phos), ribosomes, the mitochondrial unfolded protein response (mitoUPR), and “Determination of lifespan” were the most enriched terms in the insoluble proteome (Fig.1 D, Supp. Table 2). 42 of the 70 proteins in the *C. elegans* proteome bearing the annotation “Determination of Lifespan” became insoluble due to Aβ (Fig. 1D). Using STRING clustering, we found that these 42 proteins form a functionally related protein-protein interaction (PPI) network, connecting protein folding machineries with translation and oxidative phosphorylation (STRING, p<10-6) (Fig. 1E).

Normal aging causes a significant increase in the amount of insoluble protein and we noticed many of the same proteins that typically become insoluble during normal aging were being driven to become insoluble by Aβ^4,5,9^. This led us to question the extent of the similarity between the two insoluble proteomes. To assess this rigorously, we generated a list of proteins that robustly insolubilize during normal aging by overlapping two published aging insoluble proteomes from different laboratories^5,29^. This resulted in an overlap of 457 proteins that reliably become insoluble during aging (p <0.0001, Fischer’s exact test) (Fig. 1G). When we compared this with the Aβ insoluble proteome, we found that 66%, or 305 proteins, became insoluble under both conditions (p < 0.0001, Fischer’s exact test) (Fig. 1G). Moreover, when we compared the GO Biological Processes (BPs) represented in the aging and Aβ insoluble proteomes, we uncovered an 89% overlap, indicating that aging and Aβ drive insolubility of proteins involved in almost identical biological processes (Fig. 1H). The fact that two thirds of proteins that become insoluble during normal aging also do so under Aβ expression suggested to us that we identified a core set of vulnerable proteins that become insoluble under stress conditions, which we refer to herein as the ‘Core Insoluble Proteome’ (CIP).

We suspected from previous work^4^ that the CIP might be particularly enriched with regulators of aging. Indeed, we found that the CIP was highly enriched with modulators of lifespan. Specifically, 100 proteins, or roughly one third of the CIP, modulate lifespan according to the GenAge database (Fig.2A)^62^; with 80% of those proteins extending lifespan when their expression is reduced or eliminated (Fig. 2B). Furthermore, using published whole-genome RNA interference (RNAi) screens^40–44^, we found that knock down of approximately one in six of the proteins in the CIP has been shown to improve disease pathology across Huntington Disease (HD) and Parkinson Disease (PD) proteinopathy models, suggesting that the CIP might be enriched with regulators of neurodegeneration (Supp Table 5). This led us to directly question if the orthologues of CIP proteins had been implicated in neurodegenerative proteinopathies in humans. To do so, we performed disease pathway enrichment analysis with the human orthologues of the CIP and found a highly significant enrichment for a range of neurodegenerative diseases including prion disease (Pr), AD, PD and HD (Fig. 2C, Supp. Tables 6 & 7). In almost all case, the proteins were annotated against all four neurodegenerative diseases, suggesting a common mechanism might underly their associations with disease (Supp. Tables 6 & 7). These data point towards protein insolubility as the common mechanism.

**Figure 2:**
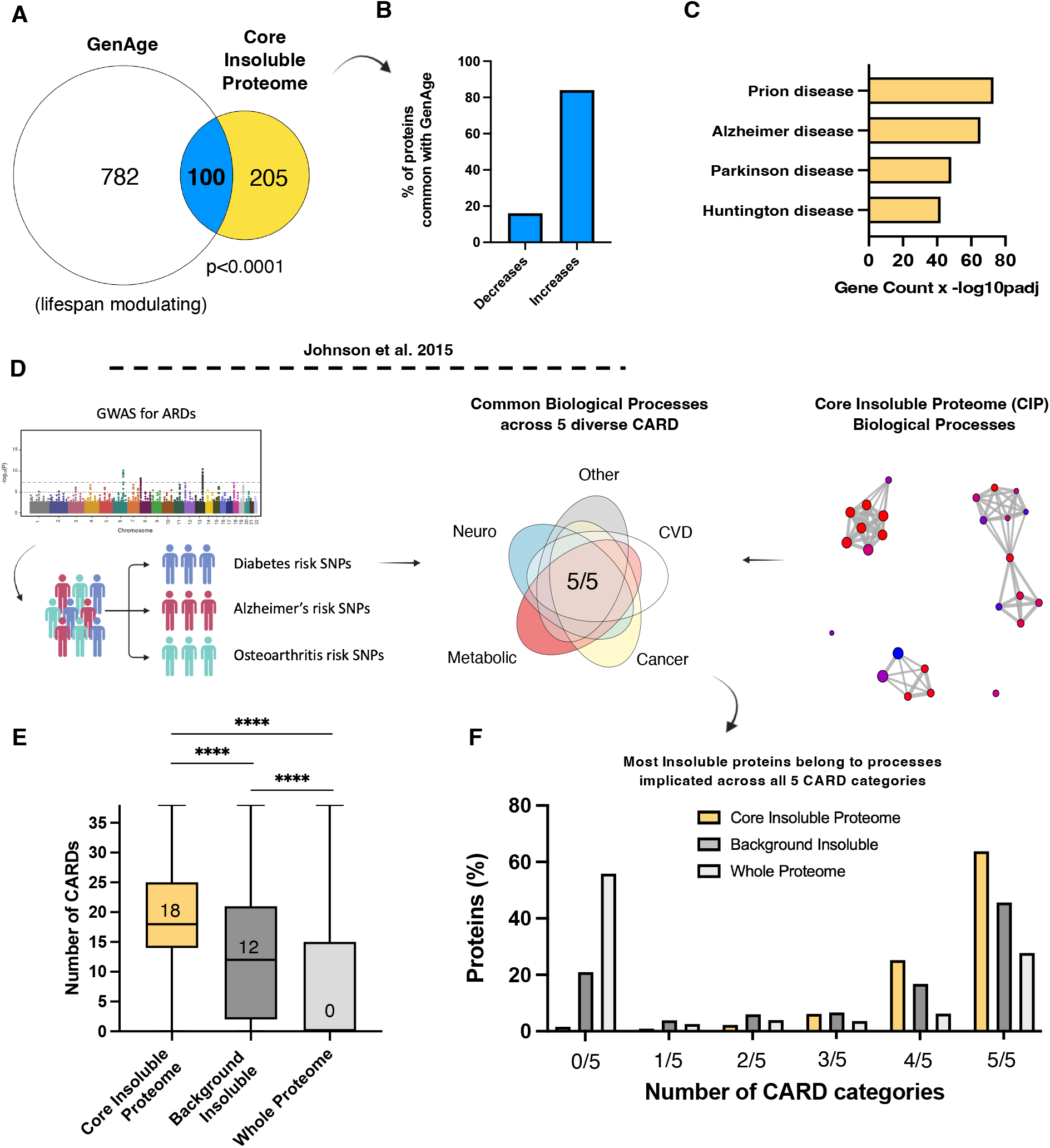
The Core Insoluble Proteome is enriched with modulators of aging and age-related disease. **A**. Overlap of lifespan modulating proteins from GenAge database with the core insoluble proteome, p<0.0001 Fischer’s Exact Test **B**. Barplot summary of published lifespan experiments where the 100 overlapping genes were either mutated or knocked down. **C**. Enriched GO terms for neurodegenerative diseases using human orthologs of CIP proteins, Benjamini Hochberg, FDR <1%. **D**. Schematic representation of GWAS analysis used to compare GO Biological Processes (BP) shared across chronic, age-related diseases (CARDs) with the BPs represented in the CIP **E**. Distribution of the number of CARDs that share biological processes with proteins found in CIP, the background insoluble proteome, and the C. elegans reference proteome with the median value displayed, Mann Whitney U test **F**. Frequency distribution of the number of broad CARD classes that share one or more biological processes with proteins found in the CIP, the insoluble proteome, and the proteome.

### Protein insolubility impacts biological processes that increase human age-related disease risk

In addition to bearing annotations for neurodegenerative disease pathways, we observed that many insoluble proteins bore pathway annotations for non-neurological, age-related diseases, such as non-alcoholic fatty liver disease and diabetic cardiomyopathy (Supp. Table 8A & 8B). This led us to question if the relationship between the insoluble proteome and chronic age-related disease (CARD) could go beyond simply neurodegeneration. Previous GWAS analyses demonstrated that common biological processes underly a whole array of diverse CARDs^45^. We therefore applied this analysis to query the commonality between biological processes impacted by protein insolubility and biological processes implicated in CARD risk^45^ (Fig. 2D). We compared biological processes against those implicated in risk of 38 distinct CARDs. We found that the average protein in the CIP was found to share biological processes with 18 of the 38 different CARDs (Fig. 2E). Moreover, 88% of the CIP proteins shared annotations with diseases spanning four or more of the following five CARD categories: neurodegenerative, metabolic, cancer, cardiovascular, and ‘other’ (which captured disparate CARDs such as macular degeneration, rheumatoid arthritis, and osteoporosis) (Fig. 2E&F). These associations were enriched compared with the experimental background insoluble proteome (ie. all proteins that could be identified in the insoluble fraction across all experimental conditions) and highly enriched compared with proteins in the proteome as a whole, in which the average protein shared processes with 0 CARDs (Fig 2E).

It is worth noting that TOMM40, which we identified in the Aβ-driven insoluble proteome and two independent aging-driven insoluble proteome datasets^10,29^, is one of only a very small fraction of SNPs (2.5%) that increase disease risk for more than 3 broad classes of CARD suggesting that TOMM40 insolubility could be a central driver of age-related disease^45^.

The biological processes common to the insoluble proteome and all 5 broad CARD categories included: immune activation and stress response pathways; growth signaling, such as fibroblast growth factor (FGF) and WNT signaling; RNA splicing and regulation of expression; and tissue homeostatic processes important for development and wound healing, such as ECM organization, cell migration and cell differentiation (Fig. S2A). Except for a few rare cases, soluble proteins lose their biological function when they become insoluble and therefore these data provide suggestive evidence that insolubility of a core set of vulnerable proteins might promote risk of not only neurodegenerative disease but CARDs more broadly.

### The CIP can be used to identify therapeutic targets

Given that one in six of the CIP proteins has been shown to alleviate toxicity across HD and PD models, we speculated that these insoluble proteins could be playing a direct role in the toxicity of Aβ, and that reducing the expression of the insoluble proteins might be beneficial. To test this, we knocked down the expression of CIP-encoding genes using RNAi and measured paralysis. The same experiment was performed with wild-type worms to rule out any non-specific effects on muscle function. We found that, of the 23 genes knocked down, 12 significantly impacted disease pathology, eight had no impact, and three were either lethal or significantly delayed development and so could not be tested. Therefore, of the CIP genes we were able to test, 60% significantly impacted Aβ toxicity (Fig. 3A), Contrary to our initial hypothesis, however, most of these genes (7/12) resulted in exacerbation of paralysis rather than protection; suggesting that knocking down expression of some proteins could be further exacerbating their loss of function caused by insolubility. When compared against a knockdown screen of ∼8000 protein-coding genes in a similar muscle-expression Aβ worm model, our results represent a roughly 60-fold greater hit rate, suggesting that the CIP is highly enriched with disease-modifying proteins^46^.

**Figure 3:**
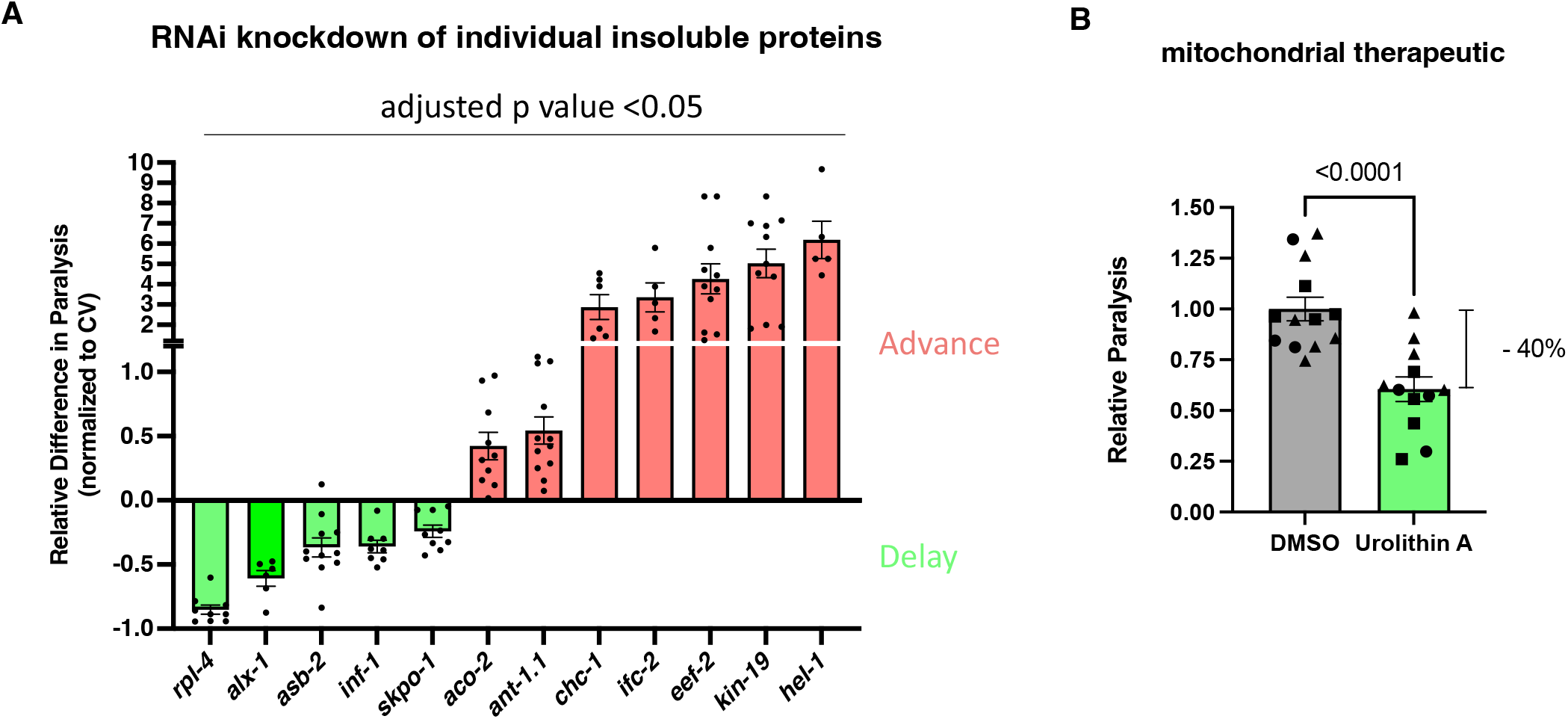
Targeting the insoluble proteome genetically or pharmacologically modulates Aβ toxicity. **A**. Statistically significant hits from RNAi paralysis screen against CIP proteins. Each point represents the relative proportion of paralyzed worms on that plate normalized to the average paralysis on control vector for that experiment, approx. 40 worms per plate, 2-3 separate experiments, error bars = SEM, for statistical tests see Supplementary Table 10A, all p<0.05 with Bonferroni correction. **B**. Barplot of paralysis under 50μM UA treatment, 40 worms per plate, 3 experiment replicates, replicate experiments are indicated by shape, error bars = SEM, unpaired t test.

Since we observed strong evidence of mitochondrial protein unfolding in the insoluble proteome, we hypothesized that directly targeting mitochondrial protein homeostasis with a small molecule might represent a good strategy to ameliorate amyloid toxicity. Urolithin A (UA) is a mitophagy-inducing, natural product derived from the bacterial metabolism of foods containing ellagitannins, such as pomegranate seeds and walnuts, in the gastrointestinal tract^47–49^. UA increases lifespan and protects neurons from amyloid toxicity in worms through a mitophagy-dependent mechanism^50,51^. UA supplementation in older adults was shown to improve muscle function in a phase II clinical trial for age-related frailty^52,53^. We treated Aβ-expressing worms with UA for 18 hours prior to inducing Aβ expression and then scored paralysis after 24 hours. We measured a robust and significant decrease in the percentage of paralyzed worms at 24 hours (Fig. 3B). Taken with recent Aβ mouse model data, this supports that targeting mitochondrial protein homeostasis by inducing mitophagy represents a reasonable strategy for preventing or treating Aβ toxicity^54^.

## Discussion

In this study we used an unbiased proteomics approach to assess the impact of Aβ on protein insolubility. We discovered that Aβ expression is sufficient to cause proteome-wide protein insolubility and in particular unfolding of the mitochondrial ETC and membrane transporters. We found that the Aβ-driven insoluble proteome is highly similar to the aging-driven insoluble proteome, suggesting there exists a core sub-proteome, vulnerable to insolubility under stress conditions. This core insoluble proteome was remarkably enriched for regulators of lifespan and disease pathology in *C. elegans*; with a particularly strong enrichment for regulators of neurodegenerative proteinopathies. Using analysis of GWAS data^45^ we uncovered suggestive evidence that protein insolubility could link diverse CARDs by the disruption of common biological processes. This finding is compelling considering the geroscience hypothesis, which “posits that aging physiology plays a major role in many — if not all — chronic diseases”^55^. While purely correlative at this stage, it leads us to hypothesize that insolubility could be a contributing factor to the etiology of CARDs more generally. While this study focused on the impact of Aβ on protein insolubility we might expect that other aggregation-prone proteins that disrupt protein folding mechanism such as polyglutamine repeat proteins or α-synuclein could have a similar impact on overall protein solubility which warrants further study.

The Aβ-driven insoluble proteome data is consistent with evidence in yeast and human cells that Aβ, and specifically the Aβ_1–42_ peptide, interferes with the import of mitochondrial protein precursors into the mitochondria by directly interacting with the TOM proteins^56–59^. The present study provides the first evidence that outer membrane proteins themselves become insoluble due to Aβ, including GOP-3 (SAMM50) of the SAM complex which has not been implicated before. This is consistent with our observation that the insoluble proteome is highly enriched with nuclear-encoded but not mitochondrial-encoded ETC complex proteins. Further, we found that targeting mitochondrial protein homeostasis with an established mitophagy-inducing compound, Urolithin A, was efficacious in preventing Aβ toxicity which may explain its observed therapeutic effects in Aβ-expressing pre-clinical mouse models^54^. Interestingly, while many interventions that increase lifespan in *C. elegans* are associated with reduced fertility, UA was shown to increase reproductive span by increasing mitochondrial quality in the oocyte, further supporting its potential as a gerotherapeutic^60^. These data clearly support the notion that Aβ disrupts mitochondrial function through loss of mitochondrial protein homeostasis and possibly mitochondrial outer membrane protein transporters which warrants further attention in the development of AD therapeutics.

Here we demonstrated that insoluble proteomics can be used to identify modifiers of Aβ toxicity. We show that reducing the levels of specific proteins in the insoluble proteome can either ameliorate or exacerbate Aβ toxicity in *C. elegans*. We observed counter-intuitive effects which could point towards specific disease-modifying mechanisms for future research. Firstly, knock down of the large ribosomal protein RPL4 caused a robust suppression of paralysis. This was anticipated because we assumed this would reduce global translation and thus reduce the overall demand on the proteostasis machinery. However, knock down of the translation elongation factor EEF-2, which should have similarly decreased global translation, had the opposite effect, resulting in a profound increase in paralysis. Similarly, we also observed a counter-intuitive result when targeting insoluble mitochondrial proteins. Knocking down the mitochondrial ATP synthase peripheral stalk protein, ASB-2, for example was found to significantly delay paralysis. In contrast, knocking down the mitochondrial membrane transporter ANT-1.1 or the aconitase enzyme ACO-2 led to significantly worse paralysis. RNAi against ASB-2 and other subunits of the ATP synthase complex are known to increase lifespan^61,62^, possibly through a mito-hormetic response, and this could explain the protective effects observed in our model. However, ANT-1.1 and ACO-2 knock down have also been shown to increase lifespan by several independent groups^4,62–65^, demonstrating a more complex picture. While these data could point to a compensation response, we speculate that certain insoluble proteins might serve a cytoprotective role in sequestering misfolded proteins or, in this case, toxic Aβ monomers, into inert, insoluble aggregates. Knock down of other members of the insoluble proteome might initiate a protective response at the level of transcription or protein homeostasis machinery.

Over 50 biological processes with known links with age-related disease risk were shown here to be linked with insolubility during normal aging (Fig S2). These data suggest that clinicians and scientists working on CARD mechanisms could benefit from considering protein insolubility when studying the interplay between aging and their disease of interest; particularly of proteins implicated in this study. We propose that evidence exists to support this approach in thinking about therapeutic strategies for age-related diseases; for example, the small molecule, HBX, which slows aging by increasing protein homeostasis in invertebrate models had unexpected, positive effects on bone health during aging in the laboratory mouse^66^. Based on the discovery of a core insoluble sub proteome and the processes implicated in CARD risk, targeting the insoluble proteome provides an encompassing strategy for the prevention and treatment of disparate age-related diseases.

## Experimental Methods

### Animal Strains

The following strains were used to generate the proteomics data: GMC101 (dvIs100 [*unc-54*p::A-beta-1-42::*unc-54* 3’-UTR + *mtl-2p*::GFP]), CL2122 (dvIs15 [(pPD30.38) *unc-54*(vector) + (pCL26) *mtl-2*::GFP]). GMC101 was used to test Urolithin A for protection against paralysis.

The following *C. elegans* strain was generated in for this study, GL399 (dvIs100 [*unc-54*p::A-beta-1-42::*unc-54* 3’-UTR; *spe-9*(hc88) I; *rrf-3*(b26) II.), by crossing TJ1060(*spe-9*(hc88) I; *rrf-3*(b26) II) with GMC101 (dvIs100 [*unc-54p*::A-beta-1-42::*unc-54* 3’-UTR + *mtl-2p*::GFP]). GL399 was used for all RNAi paralysis assays.

### Animal Maintenance

GMC101 and CL2122 worms were maintained at 20ºC on 60mm Nematode Growth Media (NGM) agar plates seeded with OP50 *E. coli*. Worms were maintained by transferring 30-50 eggs to a fresh plate on Monday and Friday. GL399 was maintained at 15ºC on 60mm NGM agar plates seeded with OP50 *E. coli*. Worms were maintained by transferring 30-50 eggs to a fresh plate weekly.

### Urolithin A Paralysis Assay

A 20mM stock of Urolithin A (UA) was prepared in sterile DMSO and stored in aliquots at -20ºC. From the stock solution, 130μL of the working solution (50μM) was prepared by mixing 7.5μL of stock solution (or DMSO only for control plates) with 125.5μL of distilled sterile water and was added to the top 35mm NGM plates (3 mL NGM agar) pre-seeded with OP50 *E. coli*. A population of synchronized GMC101 worms was generated by allowing Day 2 adult worms to lay eggs for 2 hours at 20ºC on 60mm NGM agar plates seeded with OP50. Progeny were allowed to develop to larval stage 4 (L4) and then transferred to NGM plates overlaid with 50μM UA (or DMSO only for control plates, 0.05% DMSO). After 18 hours, these plates were moved to 25ºC to initiate Aβ expression. Worms were scored for paralysis after a further 24 hours. To score paralysis in an unbiased way, worms were moved to one quadrant of the plate and scored as paralyzed if they were unable to moved away.

### RNA Interference Paralysis Assay

When the GMC101 strain is cultured on the RNAi HT115 bacteria, paralysis is significantly delayed. In addition, sterilization via FUDR treatment completely protects worms from Aβ-induced paralysis. Therefore to perform an RNAi paralysis screen in the absence of FUDR, it was necessary to generate a temperature-sensitive sterility mutant expressing Aβ. We therefore generated a new strain, GL399, by crossing the GMC101 Aβ expressing strain with TJ1060 which possesses a temperature-sensitive *Spe-9* mutation. Consequently, when GL399 is cultured at 25ºC, worms become infertile at the same time as expressing Aβ. This allowed us to perform the paralysis assay over a longer time course. *Spe-9* has no impact on lifespan and did not prevent paralysis in response to Aβ expression. A population of synchronized GL399 worms was generated by allowing Day 2 adult worms to lay eggs for 2 hours at 20ºC on 60mm NGM agar plates seeded with HT115 *E. coli* expressing the scrambled interfering RNA, or control vector RNAi. Progeny were allowed to develop to for 48 hours and then transferred to NGM plates seeded HT115 *E. coli* expressing either the scrambled control vector RNAi or RNAi against the gene of interest, and immediately shifted to 25ºC to initiate Aβ expression. Worms were scored for paralysis after 50 hours and 72 hours in order to identify RNAi conditions that advance or delay paralysis, respectively. To score paralysis in an unbiased way, worms were moved to one quadrant of the plate and scored as paralyzed if they were unable to moved away. Graphed results reflect the summarized findings of the two timepoints. Statistical tests performed and significance values are contained within Supplementary Table 10A.

### Insoluble Protein Extraction

Isolation of SDS-insoluble proteins from worms was completed as described in detail in^29^. Briefly, 200 day 2 adult worms were allowed to lay eggs for 5 hours on 100mm NGM agar plates seeded with 4x concentrated OP50 *E*.*coli*. After 50-52 hours worms were collected in sterile S-basal solution (5.85 g NaCl, 1 g K_2_ HPO_4_, 6 g KH_2_PO_4_, 1 ml cholesterol (5 mg/ml in ethanol) per 1L sterile H_2_O) and transferred to fresh 4x OP50 seeded plates. At 72 hours post egglay worm plates were transferred to 25ºC to initiate Aβ expression. 24h hours later worms were collected in sterile S-basal. The worms were allowed to settle under gravity and the supernatant solution was removed. This step ensured larval worms would not be taken forward for insoluble protein isolation. The worms were washed several times to remove all bacteria and larvae and then the supernatant was removed and the tubes flash-frozen in dry ice/ethanol.

Worm pellets were thawed in the presence of detergent-free lysis buffer containing protease inhibitor (20 mM Tris base, pH 7.4, 100 mM NaCl, 1 mM MgCl_2_). Thawed pellets were then sonicated in a water bath kept at 4ºC on maximum intensity for 30 s ON and 30 s OFF to prevent sample overheating. After the first round of sonication for 10 cycles (10 min) pellets were checked under the light microscope to ensure worms were efficiently lysed. Lysates were clarified by centrifugation at 3,000g for 4 minutes. Supernatant was collected and total protein quantified by BCA.

Insoluble proteins were extracted by aliquoting lysate equivalent to 2mg of total protein and spinning at 20,000g for 15 minutes at 4ºC. The supernatant was collected as the aqueous-soluble fraction. The insoluble protein pellet was resolubilized in 1%(w/v) SDS lysis buffer (20 mM Tris base, pH 7.4, 100 mM NaCl, 1 mM MgCl_2_ plus protease inhibitor) and centrifuged at 20,000g for 15 minutes at room temperature. The supernatant was collected as the 1% SDS-soluble fraction. The pellet was then resolubilized and washed twice more in 1% SDS lysis buffer and the supernatant fractions kept. The remaining SDS-insoluble pellet was resolubilized in sterile 70% (v/v) formic acid and sonication in a water bath for 30 minutes. Samples were then dried in a vacuum concentrator for 1 hour to completely remove formic acid. 1 x LDS sample gel buffer was added to the dried pellet and the samples were heated to 95ºC for 10 minutes. Samples were briefly centrifuged to collect condensation and stored at -80ºC or run onto a 4-12% NUPAGE Bis-Tris gel.

### Mass Spectrometric Acquisitions

Dried pellets were dissolved in 40 μL of 1x LDS sample gel buffer, incubated at 95 °C for 10 min, vortexed, and spun down. Solubilized samples were run in pre-cast NuPAGE 4-12% gradient acrylamide Bis-Tris protein gels (Invitrogen) for 20 minutes to concentrate the proteins in a single band in the stacking gel. For in-gel digestion, the gel bands were diced, collected in tubes, and dehydrated with a dehydration buffer (25 mM ammonium bicarbonate in 50% acetonitrile (ACN) and water). The gel samples were dried in a vacuum concentrator, reduced with 10 mM dithiothreitol (DTT) in 25 mM ammonium bicarbonate (pH 7-8) and incubated for 1 hour at 56 °C with agitation, then alkylated with 55 mM iodoacetamide (IAA) in 25 mM ammonium bicarbonate (pH 7-8), and incubated for 45 minutes at room temperature in the dark. The diced gel pieces were washed with 25 mM ammonium bicarbonate in water (pH 7-8), dehydrated again with the dehydration buffer, and dried in a vacuum concentrator. Afterwards, 250 ng of sequencing-grade trypsin in 25 mM ammonium bicarbonate (pH 7-8) was added to each sample. Gel pieces were vortexed for 10 min, briefly spun, and incubated at 4 °C for 30 min without mixing. Gel pieces were covered with 25 mM ammonium bicarbonate (pH 7-8) and incubated overnight for 16-20 h at 37 °C with agitation. Subsequently, the digested peptides were further extracted, as gel pieces were subjected to water and then two separate additions of a solution containing 50% ACN, 5% formic acid (FA) in water. After each addition of solution, the sample was mixed for 10 minutes, and then the aqueous digests from each sample were transferred into a new tube. These pooled peptide extractions were dried in a vacuum concentrator until completely dry. Proteolytic peptides were re-suspended in 30 μL of 0.2% FA in water and desalted using stage-tips made in-house containing a C_18_ disk, concentrated in a vacuum concentrator, and re-suspended in 15 μL of 0.2% FA in water and 1 μL of indexed Retention Time Standard (iRT, Biognosys, Schlieren, Switzerland).

Samples were then subjected to mass spectrometric analysis using a high-performance liquid chromatography (HPLC) system combined with a chip-based HPLC system (Eksigent nano-LC) directly connected to a quadrupole time-of-flight mass spectrometer (TripleTOF 5600, a QqTOF instrument) as detailed in our previous step-by-step protocol^29^.

### DIA/SWATH Data Processing and Statistical Analysis

Each sample was acquired in data-dependent acquisition (DDA) mode to build peptide spectral libraries, as we previously described^29^. Data-Independent Acquisition (DIA)/SWATH data was processed in Spectronaut (version 12.0.20491.3.15243) using DIA. Data extraction parameters were set as dynamic and non-linear iRT calibration with precision iRT was selected. DIA data was matched against an in-house *Caenorhabditis elegans* spectral library that provides quantitative DIA assays for 5,461 control and 10,140 GMC 101 *C*. elegans peptides corresponding to 1,223 protein groups for control and 1,839 protein groups for GMC 101 and supplemented with scrambled decoys (library size fraction of 0.01), using dynamic mass tolerances and dynamic extraction windows. DIA/SWATH data was processed for relative quantification comparing peptide peak areas from different days. Identification was performed using 1% precursor and protein q-value. Quantification was based on the peak areas of extracted ion chromatograms (XICs) of 3 – 6 MS2 fragment ions, specifically b- and y-ions, with q-value sparse data filtering and iRT profiling being applied. For this sample-set, local normalization was not implemented. Differential protein expression analyses for all comparisons were performed using a paired t-test, and p-values were corrected for multiple testing, using the Storey method. Specifically, group wise testing corrections were applied to obtain q-values. Protein groups with at least two unique peptides, q-value < 0.01, and absolute Log_2_(fold-change) > 0.58 are significantly altered (Supp. Tables 1,11A,11B).

### Data Accession

Raw data and complete MS data sets have been uploaded to the Mass Spectrometry Interactive Virtual Environment (MassIVE) repository, developed by the Center for Computational Mass Spectrometry at the University of California San Diego, and can be downloaded using the following link: https://massive.ucsd.edu/ProteoSAFe/private-dataset.jsp?task=d7c68fbc04ce4f5e9c757cd14daa7585 (MassIVE ID number: MSV000092250; ProteomeXchange ID: PXD043250). Enter the username and password in the upper right corner of the page: Username: MSV000092250_reviewer; Password: winter

### Computational Methods

Enrichment analysis: All enrichment analyses were performed using significantly increased proteins (Log_2_(fold-change) > 0.58, Q < 0.01) and gProfiler^67^ with Benjamini-Hochberg <5% FDR correction. Enrichment results are presented as the product of gene count and FDR-corrected p value. Human orthologs were derived from Ortholist^36^.

GWAS analysis: All SNP GO biological process data for the present study was taken from Johnson et al.^45^. All GO biological process data for insoluble proteins was downloaded from DAVID^68,69^ and Uniprot^70^. For comparison between GO biological processes enriched in CARD risk with those represented in the insoluble proteome and CIP (the overlap between the Aβ-driven and aging-driven insoluble proteome), we determined overlaps between GO term annotations for the proteins and the CARDs. We used these overlaps then to determine the shared biological processes in the CIP and insoluble proteome with the five broad age-related disease categories as defined in Johnson et al.^45^. The “background insoluble” proteome was defined as any protein which could be reliably identified in the insoluble fraction in either of the experiments ie. the experimental background. For comparison with the “whole proteome” writ large, we generated a random list of 1600 proteins from the *C. elegans* proteome using a random number generator and downloaded their GO biological process data from DAVID and Uniprot. GO biological processes shared between the CIP and all 5 diverse categories of CARD were processed through ReviGO^71^ to remove any redundant terms and create a relationship network between the processes, which we subsequently modified in Cytoscape^72^ and presented in Fig.S2.

## Acknowledgments

GJL, BS & JKA disclose support for the research of this work from NIA RF1AG057358 and GJL discloses support for the research of this work from The Larry L. Hillblom Medical Foundation Lithgow LLHF Ctr Supp. and NIA U01AG045844. BS acknowledges support of instrumentation for the TripleTOF 6600 from the NIH shared instrumentation grant 1S10 OD016281 (Buck Institute).

Graphical figures created with biorender.com

Special thanks to Dr Simon Johnson for providing source data from^45^.

## COI Statement

The authors declare no conflict of interest.

---

**Supplementary Figure 1.**
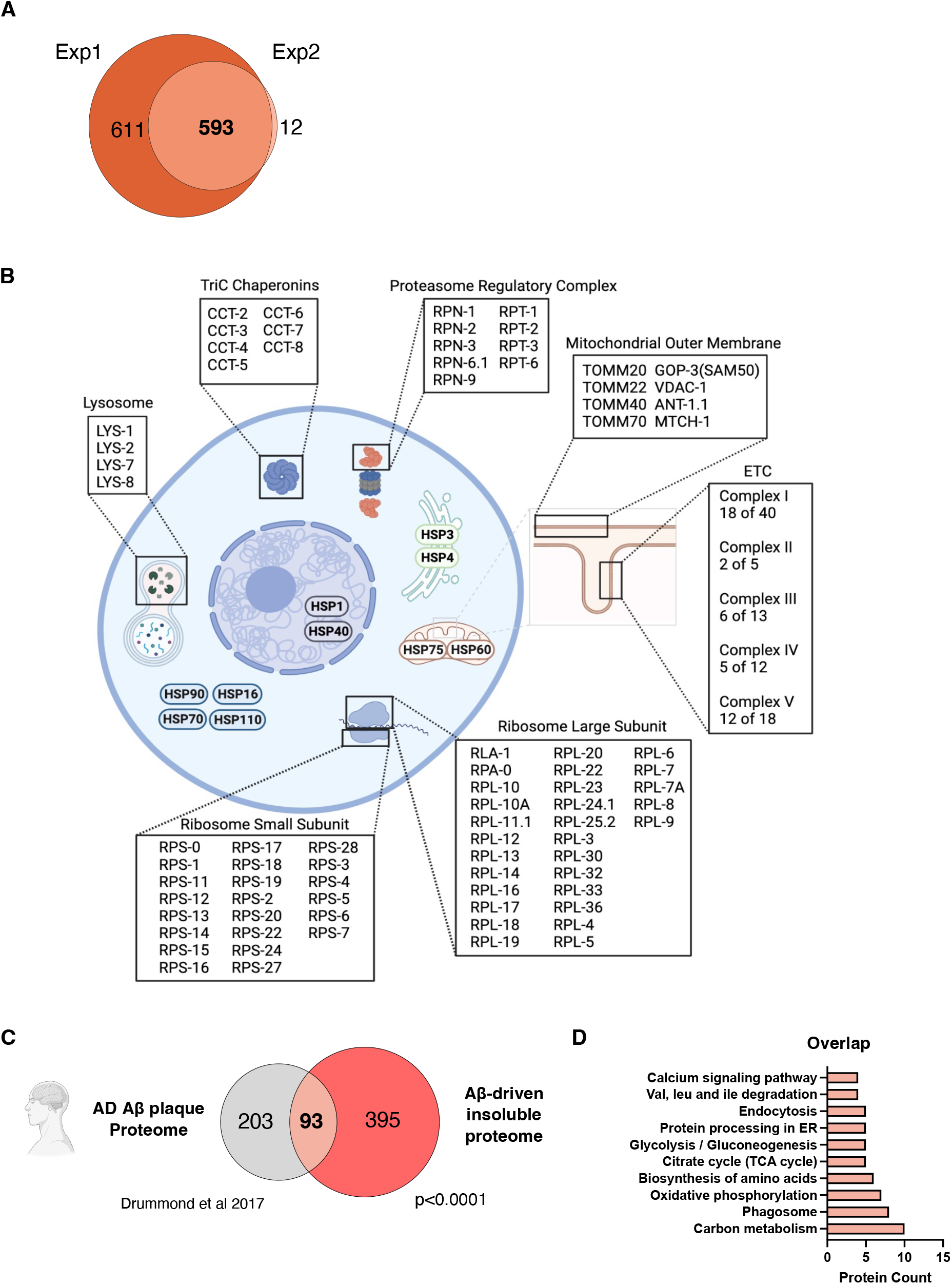
**A**. Overlap of proteins which increased in abundance in the insoluble proteome after Aβ expression. **B**. Schematic overview of proteostasis and mitochondrial proteins enriched in the Aβ-driven insoluble proteome. **C**. Overlap of proteins reliably identified in Aβ-rich plaques from AD patient brains and the Aβ-driven insoluble proteome, Fischer’s exact test. **D**. Top 10 KEGG annotations by protein count for overlapping proteins between AD senile plaque proteome (Drummond et al. 2017) and the Aβ-driven insoluble proteome (this manuscript).

**Supplementary Figure 2.**
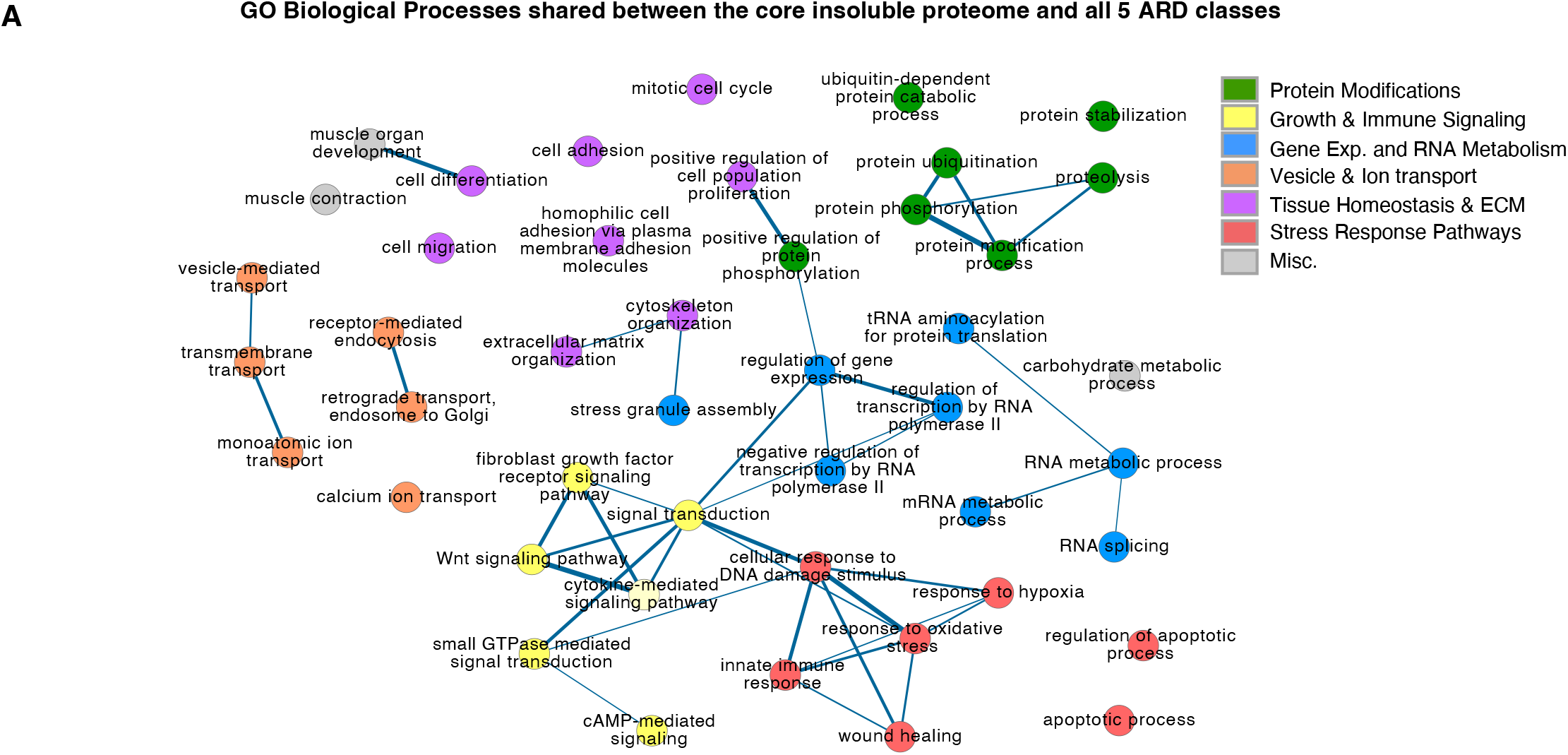
**A**. ReviGo network representation of the non-redundant GO biological processes shared between the CIP and all 5 diverse CARD categories.

## Notes

https://massive.ucsd.edu/ProteoSAFe/private-dataset.jsp?task=d7c68fbc04ce4f5e9c757cd14daa7585

### Competing Interest Statement

The authors have declared no competing interest.

### Summary of Updates

Figure 2D explanatory graphic revised to improve clarity. Supplementary figure 1C&D added for comparison with AD plaque proteome. Introduction lexical clarity improved.

